# A deep dive into the ancestral chromosome number and genome size of flowering plants

**DOI:** 10.1101/2020.01.05.893859

**Authors:** Angelino Carta, Gianni Bedini, Lorenzo Peruzzi

**Author notes:** Author for correspondence: Angelino Carta.

## Abstract

- Chromosome number and genome variation in flowering plants has stimulated a blossoming number of speculations about the ancestral chromosome number of angiosperms, but estimates so far remain equivocal.
- We used a probabilistic approach to model haploid chromosome number (*n*) changes along a phylogeny embracing more than 10 thousands taxa, to reconstruct the ancestral chromosome number of the common ancestor of extant angiosperms and the most recent common ancestor for single angiosperm families. Independently, we carried out an analysis of 1C genome size evolution, including over 5 thousands taxa.
- Our inferences revealed an ancestral haploid chromosome number for angiosperms *n* = 7, a diploid status, and an ancestral 1C = 1.73 pg. For 160 families, inferred ancestral *n* are provided for the first time.
- Both descending dysploidy and polyploidy played crucial roles in chromosome number evolution. While descending dysploidy is equally distributed early and late across the phylogeny, polyploidy is detected mainly towards the tips. Similarly, also 1C genome size significantly increases (or decreases) in late-branching lineages. Therefore, no evidence exists for a clear link between ancestral chromosome numbers and ancient polyploidization events, suggesting that further insights are needed to elucidate the organization of genome packaging into chromosomes.

## Introduction

Chromosome rearrangements are a well-known evolutionary feature in eukaryotic organisms (Coghlan *et al.*, 2005), especially in plants. The remarkable diversity of flowering plants (angiosperms) has been attributed, in part, to the tremendous variation in their chromosome number (Stebbins, 1971). This variation has stimulated a blossoming number of speculations about the ancestral chromosome number of angiosperms (Ehrendorfer *et al.*, 1968; Stebbins, 1971; Walker 1972; Raven, 1975; Grant, 1981; Soltis *et al.*, 2005), but estimates so far remain equivocal.

Each eukaryotic organism has a characteristic chromosome complement, its karyotype, which represents the highest level of structural and functional organization of the nuclear genome (Stace, 2000). Karyotype constancy ensures the transfer of the same genetic material to the next generation, while karyotype variation provides genetic support to ecological differentiation and adaptation (Stebbins, 1971, Stace, 2000). Cytogenetic studies have shown that the tremendous inter- and intra-taxonomic variation of chromosome number documented in flowering plants (Levitzky, 1931; Stebbins, 1971; Raven, 1975) is mostly driven by two major mechanisms: a) increases through polyploidy (which may entail a Whole Genome Duplication [WGD] or an increase by half of the genome, demi-duplication [Mayrose *et al.*, 2009]); b) decreases or increases through duplication or deletion of single chromosome pairs or structural chromosomal rearrangements like chromosome fusion, i.e. descending dysploidy, and chromosome fission, i.e. ascending dysploidy.

Polyploidy is a common and ongoing phenomenon, especially in plants (Wood *et al.*, 2009), that has played an important role in many lineages, with evidence of several rounds of both ancient and recent polyploidization (Jiao *et al.*, 2011; Li *et al.*, 2015; Leebens-Mack *et al.*, 2019), albeit its distribution in time remains contested (Ruprecht *et al.*, 2017). Indeed, although the crucial role of polyploidy in plant diversification on small timescales is widely accepted (Stebbins, 1971; Grant, 1981), the evolutionary significance of polyplodization for the long-term diversity of angiosperms is still controversial (Mayrose *et al.*, 2011; Soltis *et al.*, 2014; Han *et al.*, 2020). On the other hand, while dysploidy is more frequent than polyploidy in angiosperms (Grant, 1981), its adaptive consequences have been mostly unexamined (Weiss-Schneeweiss & Schneeweiss, 2013), until recent studies demonstrated its high evolutionary impact (Escudero *et al.*, 2014).

In other seed plants, Li et al. (2015) detected two WGDs in the crown node of conifer clades, namely Pinaceae and Cupressaceae, contrary to a previous study suggesting a slow and steady rate of genome size increase mainly due to the accumulation of repetitive DNA (Nystedt et al., 2013). Several rounds of polyploidy, followed by diploidization, also occurred in monilophyte ancestors (Clark et al., 2016), albeit these authors managed a limited taxonomic sampling. Diploid chromosome numbers range from 2*n* = 14 to 66 in extant gymnosperms, and from 2*n* = 18 to 1440 in monilophytes.

Chromosome number variation across angiosperm lineages spans two orders of magnitude (Stace, 2000), from 2*n* = 4 to 2*n* = 640. Previous hypotheses (Ehrendorfer *et al.*, 1968; Stebbins, 1971; Walker 1972; Raven, 1975; Grant, 1981; Soltis *et al.*, 2005) of the ancestral basic (monoploid; see Peruzzi, 2013) chromosome number *p* in angiosperms suggest low numbers, between *p* = 6 and *p* = 9. Genome size also vary tremendously across the flowering plants (Leitch *et al.*, 2005), and it is has been estimated that the ancestral genome size of angiosperms was very small (1C < 1.4 pg, Leitch *et al.*, 2005). Early hypotheses to estimate putative ancestral basic chromosome numbers placed particular attention to ‘primitive’ extant angiosperms (Raven, 1975). More recently, an ancestral chromosome number has been reconstructed using a maximum parsimony approach (Soltis *et al.*, 2005). However, although parsimony has been widely used to infer ancestral chromosome numbers, it carries significant shortcomings (Mayrose *et al.*, 2009), and more rigorous and complex models to infer chromosome number evolution are currently available (Mayrose *et al.*, 2009; Glick & Mayrose, 2014; Freyman & Höhna, 2018; Zenil-Ferguson *et al.*, 2018). Here we use probabilistic models, accounting for various types of chromosome number transitions, to reconstruct the ancestral haploid chromosome number and the occurrence of chromosome change events across the most massive data set ever assembled linking chromosome numbers to a phylogeny, sampling 10,766 taxa from 318 families (73%) and 59 orders (92%) of angiosperms. Chromosome numbers were extracted from the Chromosome Counts DataBase (Rice *et al.*, 2015), and the analyses were conducted using pruned versions of two recently published, dated mega-trees for seed plants (Smith & Brown, 2018). In addition, we explored the sensitivity of our results by conducting all analyses again using a different ultrametric tree of 1,559 taxa extracted from a recently published alternative angiosperm tree (Li *et al.*, 2019). As WGD and post-polyploid genome diploidization (PPD) are widespread across angiosperms (the latter gradually reverting the polyploid genome to one functionally diploid-like through chromosomal rearrangements, see Mandakova & Lysak, 2018), it is critical to consider changes in ploidy level besides chromosome number change dynamics. To this aim, we compared our results about chromosome number evolution with estimated rates of different shifts in ploidy level (including diploidization events, Zenil-Ferguson *et al.*, 2018) with inferred ancestral states in genome size sampling over 5,000 taxa from the Plant DNA C-values database (https://cvalues.science.kew.org/search/angiosperm). Ancestral genome size inference taken in combination with chromosome number evolution, could allow us to address genome rearrangements associated with genome size duplication across angiosperm history.

## Materials and Methods

### Phylogenetic reconstruction

We used two recently published (Smith & Brown, 2018) dated megaphylogenies for seed plants, GBMB and GBOTB, as backbones to generate two alternative phylogenies for angiosperms included in the dataset. GBMB and GBOTB were constructed using 79,874 and 79,881 taxa, respectively, available in GenBank and in a backbone provided either by Magallón et al. 2015 (GBMB) or by Open Tree of Life, version 9.1 (GBOTB). In addition, we also used a different ultrametric tree provided by a recently published plastid phylogenomic angiosperm (PPA) tree (Li *et al.*, 2019).

### Chromosome numbers collection

The haploid chromosome numbers (*n*) of the species were obtained from the Chromosome Counts Database (CCDB [Rice *et al.*, 2015]; http://ccdb.tau.ac.il/; last accessed May 2019) using the R package chrome (Pennell *et al.*, 2016). CCDB contains records from original sources that have irregularities of chromosome counts, so that the ca. 150,000 records were curated semiautomatically using the CCDBcurator package (Rivero *et al.*, 2019). After a first round of automatic cleaning, we examined results by hand using custom R scripts and corrected records where needed.

Species with unknown chromosome counts were pruned from the trees, thus we collected chromosome numbers for 10,766 taxa included in the GBMB and GBOTB phylogenetic trees, and for 1,559 taxa included in the PPA tree. In cases where multiple chromosome numbers were reported for a given taxon, the modal number was used (Mayrose *et al.*, 2009; Salman-Minkov *et al.*, 2016). For taxa with numbers suggesting different ploidy levels, we used the lowest haploid chromosome number available (Márquez-Corro *et al.*, 2019). This coding scheme allowed us to deal with the problem of the existence of different ploidy levels in a taxon and also with the low-density sampling conducted in most taxa (Márquez-Corro *et al.*, 2019). Analyses conducted using the PPA tree encountered less computation limitations, so that we were able to perform them by explicitly considering intraspecific polymorphism, allowing several chromosome numbers, together with their respective frequencies, to be set for each taxon (Glick & Mayrose, 2014).

### Genome size collection

Genome size data (1C, prime estimates) for angiosperms were taken from the Plant DNA C-values database (https://cvalues.science.kew.org/search/angiosperm, release 7.1, March 2020), managed by the Royal Botanic Gardens, Kew (Pellicer & Leitch, 2019). We gathered 5,581 and 661 taxa, respectively matching the GBMB and the PPA trees. Genome size data were log_10_ transformed to ensure the data conformed to Brownian motion evolution (Beaulieu *et al.*, 2012; Carta & Peruzzi, 2016).

### Analyses

The evolution of haploid chromosome numbers of angiosperms was inferred using chromEvol software v.2.0 (http://www.tau.ac.il/~itaymay/cp/chromEvol/index.html; Glick & Mayrose, 2014). This software determines the likelihood of a model to explain the given data along the phylogeny, based on the combination of two or more of the following parameters: dysploidization (ascending, chromosome gain rate *λ;* descending, chromosome loss rate δ), polyploidization (chromosome number duplication with rate *ρ*, demi-polyploidization or triploidization with rate *μ*) and incremental changes to the basic number *(β)* with regard to a rate of multiplication (v) that is different from a regular duplication (Mayrose *et al.*, 2009). Two additional parameters (*λ*_1_, *δ*_1_) detect linear dependency between the current haploid number and the rate of gain and loss of chromosomes. We tested 10 models based on a different combination of the parameters above. Four of these models consider only constant rates (Mc1, Mc2, Mc3, and Mc0), whereas the other four include two linear rate parameters (Ml1, Ml2, Ml3, and Ml0; Table 3). Both sets have a null model (Mc0 and Ml0) that assumes no polyploidisation events. Finally, two models (Mb1 and Mb2) consider that the evolution of chromosome number can also be influenced by the basic number and by its transition rates. The minimum chromosome number allowed in the analyses was set to 2, whereas the maximum number was set to 5 units higher than the highest chromosome number found in the empirical data. We removed all counts *n* > 43 from the analysis (5,467 counts [3,001 taxa]), because for many lineages the sampling was inadequate to reconstruct such a drastic change in chromosome number (Márquez-Corro *et al.*, 2019; Barrett *et al.*, 2019) and because of computation limitations (Zenil-Ferguson *et al.*, 2018). The branch lengths were scaled according to the software author’s instructions. The null hypothesis (no polyploidy) was tested with likelihood ratio tests using the Akaike information criterion (AIC; Burnham & Anderson, 2004). To compute the expected number of changes along each branch, as well as the ancestral haploid chromosome numbers at internal nodes, the best fitted model for both data sets was rerun using 1,000 simulations. The best model was plotted on the trees using the ChromEvol functions v0.9-1 elaborated by N. Cusimano in R.

**Table 1.**
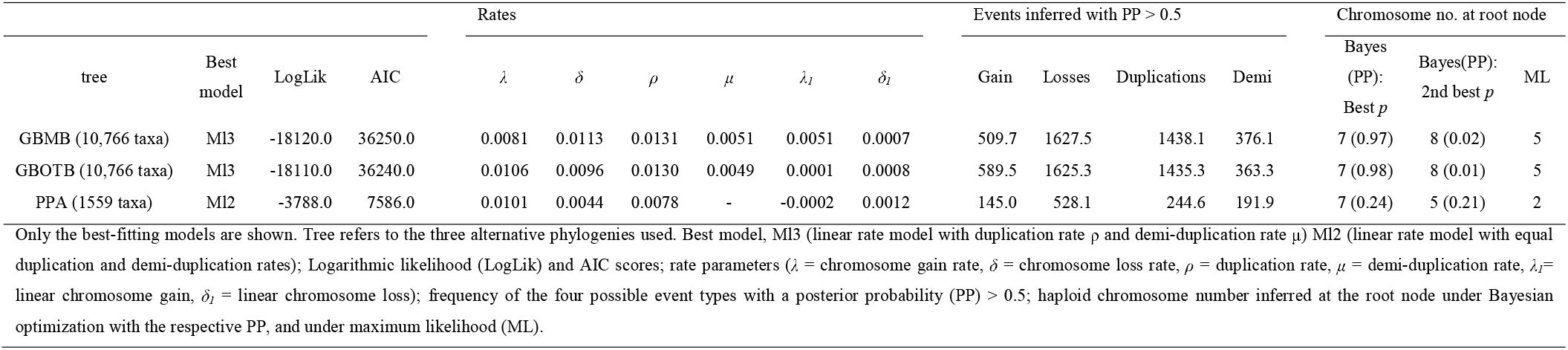
Summary of chromosome number evolutionary models and inferred ancestral haploid chromosome number (*n*) in Angiosperms under the best-fitting model.

**Table 2.**
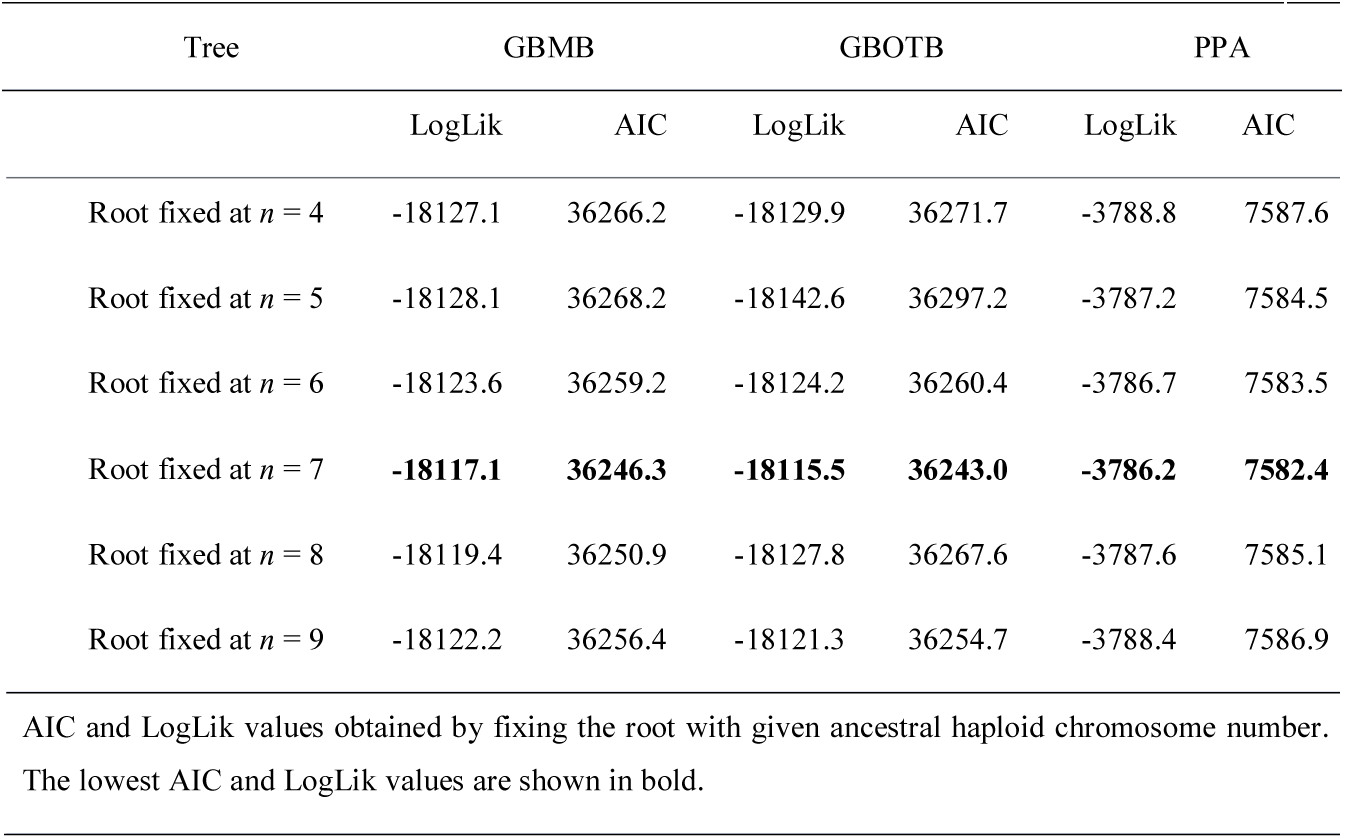
Testing hypotheses about root ancestral haploid chromosome number (*n*).

**Table 3.**
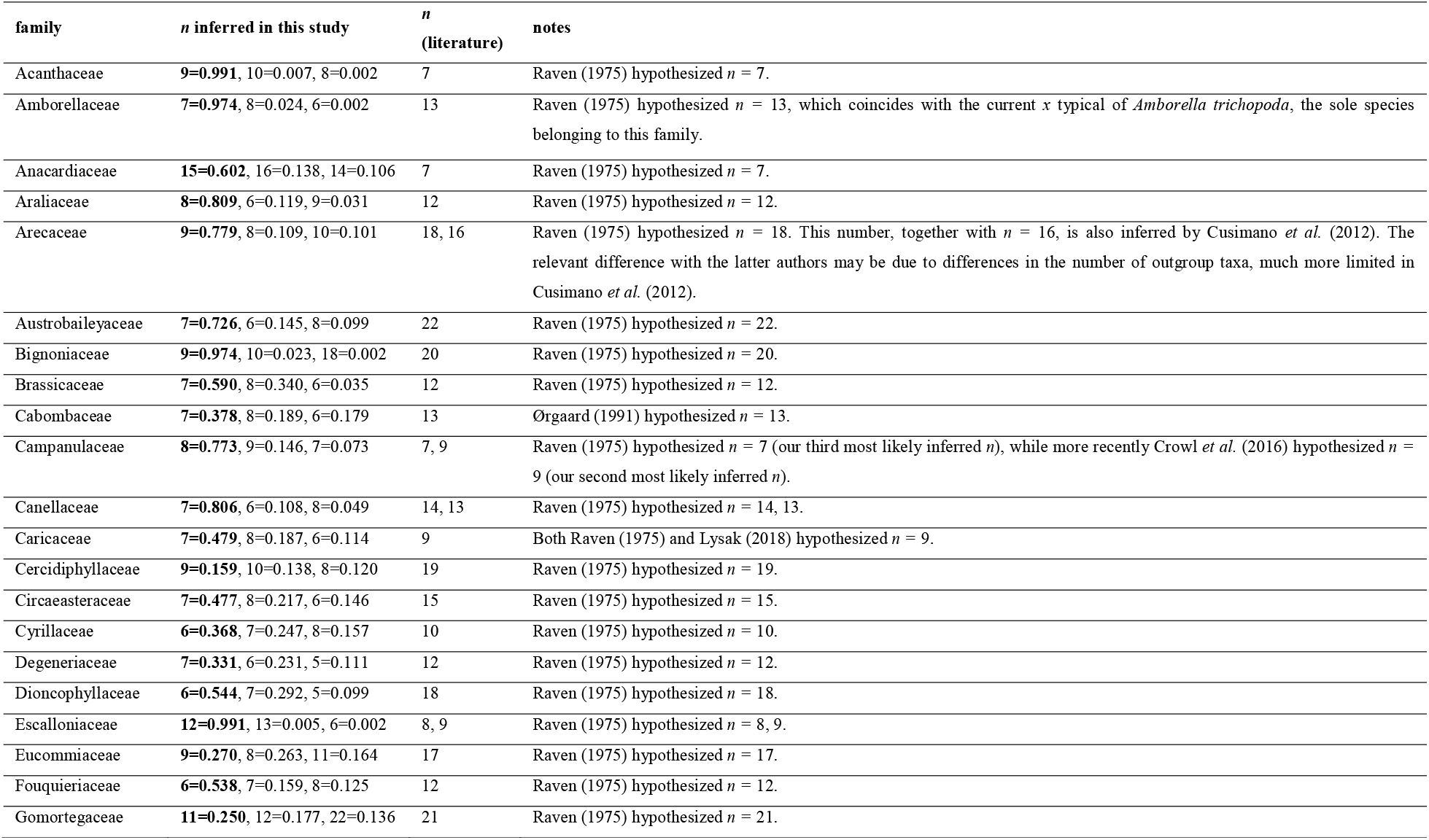
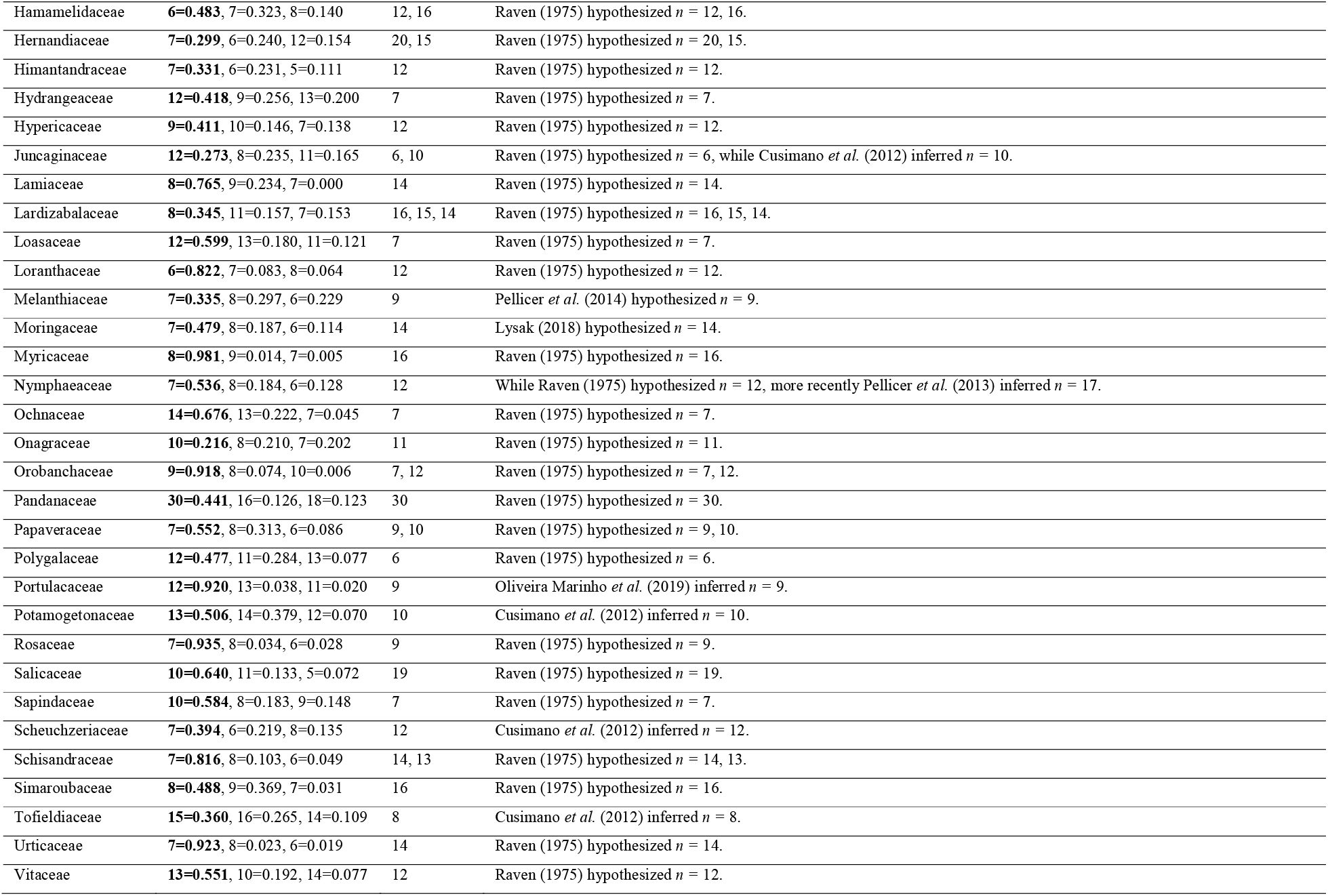
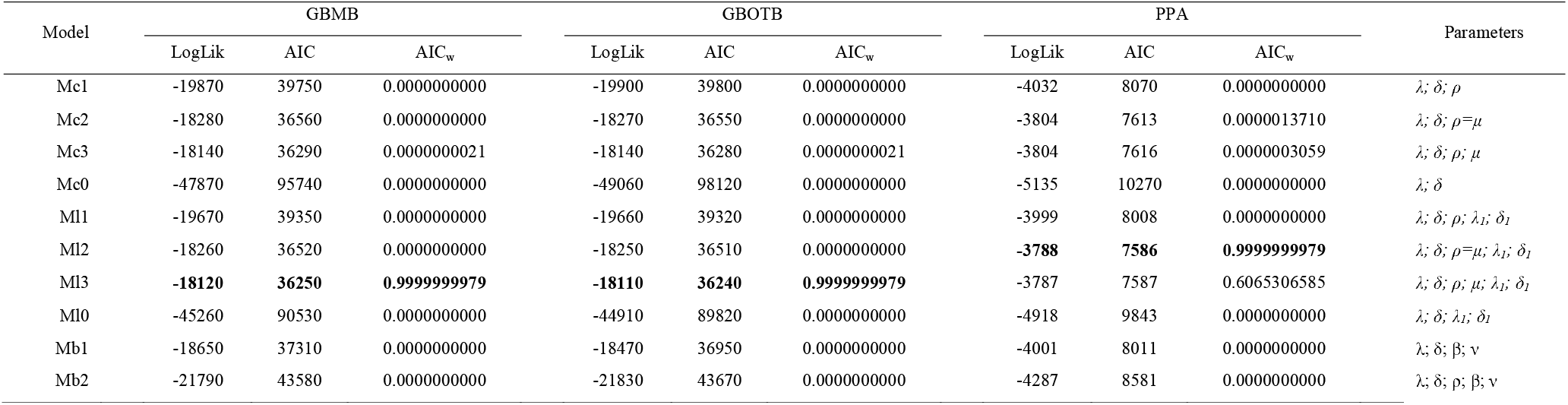
Inferred ancestral n chromosome numbers of most recent common ancestors (MRCAs) for selected angiosperm families. For each family, the three most supported numbers are given (the first most supported in boldface). Only those 52 out of 318 families for which previous - and different - hypotheses were postulated in literature are reported. For further details see Table S1.

To test which ancestral haploid chromosome number is most likely at the root of angiosperms, the following haploid chromosome numbers were fixed at the root and the likelihood of the resulting models was compared: *n* = 4,5,6,7,8,9. These numbers were tested either because considered putative ancestral character-states (Ehrendorfer *et al.*, 1968; Stebbins, 1971; Walker 1972; Raven, 1975; Grant, 1981; Soltis *et al.*, 2005), or because they were identified as the chromosome numbers showing the highest PP under our Bayesian analysis.

To further support our results, we also modelled haploid chromosome number evolution using the function Q_chromevolM3 in the R package chromploid (Zenil-Ferguson *et al.*, 2018). We then modeled ploidy change using the function PloidEvol and a subset of 3,759 taxa used in the ChromEvol analysis that have ploidy level information in https://cvalues.science.kew.org/search/angiosperm. The PlodiEvol model, implemented in the package chromploid, is a continuous time Markov chain model of ploidy level evolution, that estimates the rates of different shifts in ploidy level, including diploidization, across a phylogenetic tree, by analyzing the transitions of ploidy level among species. Optimizations were performed using a subplex algorithm (nloptr package, Ypma 2014), where the convergence criterion was based on the changes in the precision of the negative log likelihood of no more of 10^-8^. The procedure was run for at least 1,000 iterations.

Ancestral genome sizes were reconstructed using maximum likelihood and visualized on the tree with the contmap function in the R package phytools (Revell, 2012). Models of adaptive evolution at macroevolutionary scales typically rely on the Ornstein-Uhlenbeck (OU) model of trait evolution (Hansen, 1997). This model assumes that an optimum trait value identifies a selective regime acting on a lineage over the course of its history. The rate of adaptive evolution towards an optimum trait value θ is governed by α, while the constant σ^2^ describes the rate of stochastic evolution away from the optimum. Here, discrete regime shifts in genome size on the phylogeny were estimated from the data using a Bayesian reversible-jump multi-optima OU model in the R package bayou (Uyeda & Harmon, 2014). Two independent chains were run for at least 200,000 generations, with a thinning interval of 20 and the first 60,000 generations discarded as burn-in. To check the convergence of the chains we used the Gelman’s R. A half-Cauchy distribution was assigned to α and σ^2^, a normal distribution with mean and standard deviation equivalent to the distribution of values for genome size was assigned to θ, and a uniform distribution was assigned to the probability that a change in optima occurs on a given branch, with the restriction that only one change can occur on any branch. Mean genome size was estimated for each species, while within-species variation to be incorporated into the phylogenetic analysis was not estimated separately for each species, because within-species samples were limited to a few samples. OU models are particularly affected by measurement error. Hence, we estimated the pooled variance across the species and used it, weighted by sample size, to estimate the observation variance of the individual species. All analyses were performed in the high-performance computing cluster at the University of Pisa.

## Results

Regardless of the three alternative phylogenies, *n* = 7 was inferred as the ancestral haploid chromosome number with the highest posterior probability (Table 1) and likelihood (Table 2). The ancestral haploid chromosome number *n* = 7 was remarkably stable in the deepest part of the phylogeny (Fig. 1), while slight variations (± 1) in *n* were inferred at the base of some lineages. Greater variations were shown in the ancestral haploid chromosome number of many angiosperm families (see Supplementary Table S1). Monocots exhibited the largest variation of inferred *n* among Most Recent Common Ancestors (MRCAs) of plant families (Fig. 2b), paralleled by a considerable variation in current haploid chromosome numbers (Fig. 2a). Over 70% of inferred *n* in the 158 families for which previous inferences were available are in line with previous proposals. The most interesting disagreements among our inferences and those previously available in literature are reported in Table 3. For the remaining 160 families (50.3%) the first inferences are presented here.

**Figure 1.**
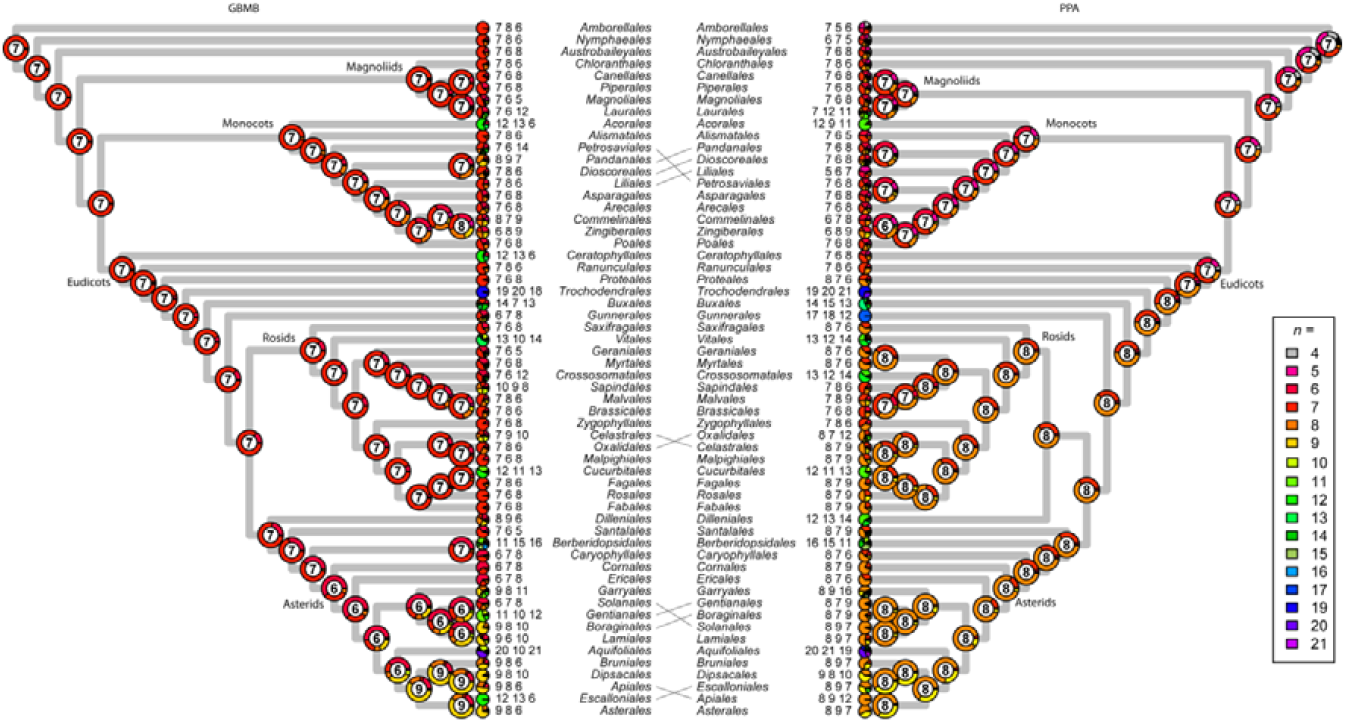
Reconstruction of ancestral haploid chromosome number (*n*) of angiosperms with the best-fitting model on the different types of trees used (GBMB and PPA). Please note that the plotted trees depict ordinal phylogenetic relationships (sub-ordinal topologies were collapsed to build the figure), and are shown without branch length information. Pie charts at nodes represent the probability of the ancestral haploid chromosome numbers inferred under Bayesian estimation; the numbers at nodes are those with the highest probability. Pie charts and numbers at the tips are the three best inferred ancestral haploid chromosome numbers per each angiosperm order. Black lines, difference in phylogenetic position between GBMB and PPA trees.

**Figure 2.**
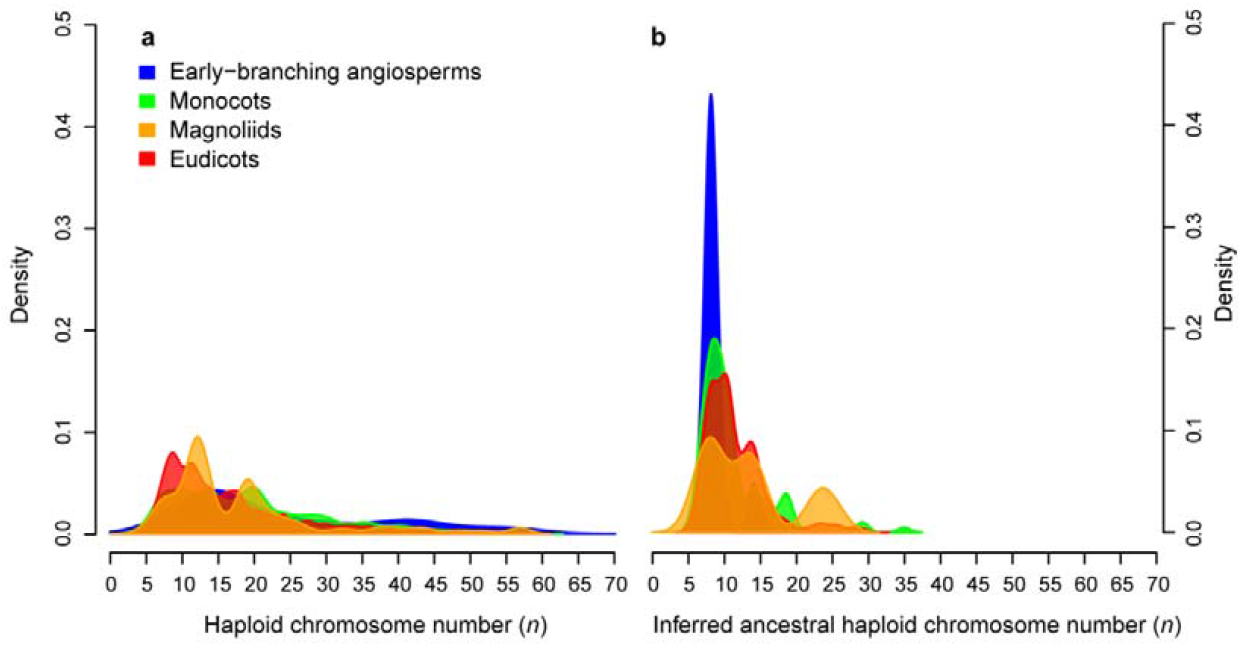
Density plots of haploid chromosome numbers (*n*) and of inferred ancestral haploid chromosome number (*n*) for each angiosperm family in four major angiosperm clades (APG IV). **a**, we identified the number of unique chromosome counts per taxon, i.e. cytotypes, from the original dataset, after excluding counts with *n* > 60, to focus on the most frequent *n* and their putative relation with inferred *p*. **b**, density plots were scaled by the Bayesian posterior probability (PP) estimated for each inferred *p*.

For both GBMB and GBOTB phylogenies, the best model considers up to six parameters (Table 4), i.e. chromosome gain, loss, duplication, demi-duplication rates and rates of gain and loss linearly dependent on the current chromosome number. Specifically, descending dysploidy, most likely through chromosome fusion, was the most common cytogenetic mechanism of chromosome number change during the evolution of flowering plants, and this is inferred both on branches leading to major clades and on terminal branches (Supplementary Figs. 1-3). Polyploidization events were also inferred in a significant number, albeit mainly on terminal branches (Supplementary Figs. 1-3). PloidEvol, having the ability to define a wider range of increasing ploidy level change events than chromosome number-based models, detected a small rate of pure duplication for even ploidy levels (*ρ* = 0.024). Instead, PloidEvol indicate that other two parameters associated with the formation of even ploidy levels are the largest: the rates of even-to-odd increase *(α* = 0.697) and of diploidization *(δ* = 0.471) and even-to-even transition (*ε* = 0.116). The stationary distribution under Ploidevol model indicates a diploid status of the root, with highest probability (0.74) followed by a tetraploid status (0.18). Analyses conducted using the PPA tree, include a lower number of sampled taxa but allowed to consider intraspecific chromosome number variation for each taxon. Nevertheless, results were consistent with those obtained using the GBMB and GBOTB phylogenies. We found indeed only minor differences at the root (Tables 1,2) and at some internal nodes (Fig. 1).

**Table 4.**
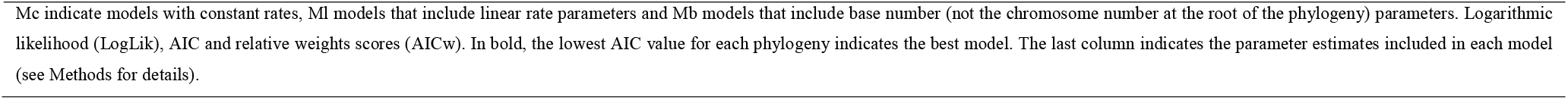
Goodness of fit of the 10 different models of chromosome number evolution applied to the three alternative phylogenies used.

The results above are also confirmed by those obtained through the chromploid software, concerning both the ancestral haploid chromosome number marginal probability (*n* = 7, 0.54; *n* = 8, 0.43; *n* = 9, 0.01) and the rate parameters, with descending dysploidy (*δ* = 0.017) and chromosome doubling (*ρ* = 0.012) largely exceeding chromosome gain (Λ. = 0.009) and demiploidization (*μ* = 0.005).

The reconstruction of genome size variation across the evolutionary history of angiosperms indicates that the ancestral holoploid genome size for angiosperms was small (1C = 1.73 pg) and mostly stable in the basal part of the tree, showing instead decreasing or increasing values along terminal branches (Fig. 3 and Figs. S4–S8). In addition, the multi-optima OU model identified several significant shifts in genome size, particularly an increase in genome size at the base of Monocots and a downsizing at base of Rosids (Figs. S6-7).

**Figure 3.**
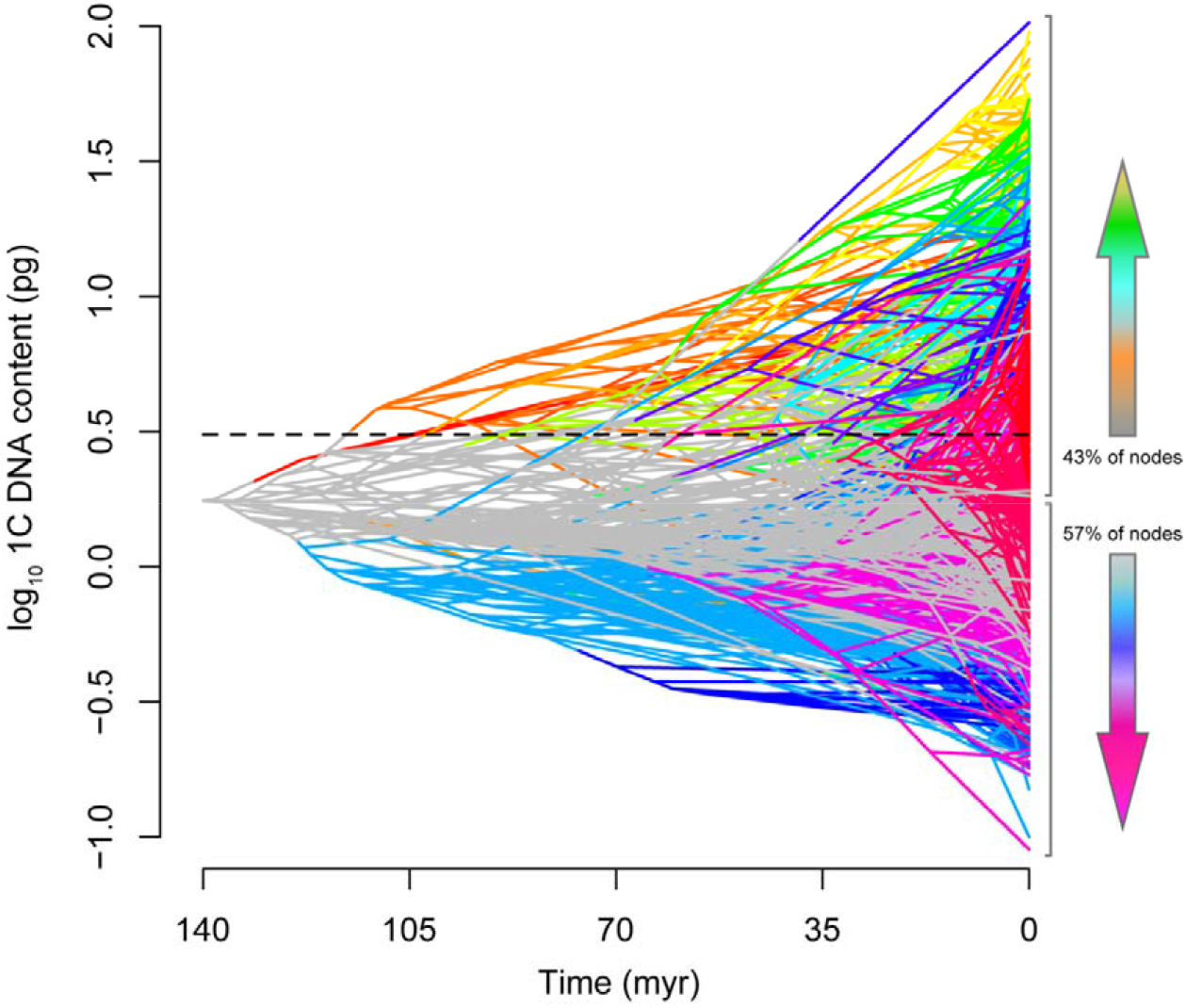
Projection of the tree into phenotype space (traitgram) for genome size evolution in angiosperms (GBMB tree). Ancestral states for the projection and discrete shifts in genome size on the phylogeny were estimated using a Bayesian reversible-jump multi-optima OU model. Branches are coloured according to their median value of theta, with the larger phenotypic values for optima being indicated by blue, green and orange while smaller optima indicated by pink. The straight dashed line coincides with a 2-fold ancestral genome size.

Inferred ancestral 1C values for angiosperm families are reported in Table S1. Overall, the majority of nodes (57%) underwent a decrease in genome size with respect to the ancestral value of 1C = 1.73 pg (0.24 in Fig. 3). In addition, a value 2-fold the ancestral one (0.48 in Fig. 3) was reached for the first time only after more than 30 million years of angiosperm evolutionary history (line in Fig. 3), in the lineage giving rise to Monocots.

## Discussion

The main results presented here were drawn using the largest dated mega-tree currently available for seed plants (but see Janssens *et al.*, 2020). Bayesian inference revealed an ancestral haploid chromosome number for angiosperms *n* = 7, reinforcing previous hypotheses (Ehrendorfer *et al.*, 1968; Stebbins, 1971; Walker 1972; Raven, 1975; Grant, 1981; Soltis *et al.*, 2005) that suggested a low ancestral basic number.

As a by-product of our study, our analyses allowed to infer *n* for single families, more than half of which are provided here for the first time, while the others are mostly congruent with previous evaluations. However, were able to highlight a number of discrepancies in ancestral chromosome numbers previously inferred for a number of families (Table 3), possibly due to traditional routine algebraic approaches, instead of phylogenetic models, to infer chromosome number changes (see Cusimano *et al.*, 2012 for further discussion), or to the analysis of more limited datasets.. For example, the inferred *n* of MRCAs of Brassicaceae, Lamiaceae, and Rosaceae are respectively 7, 8, and 7 in our study, but were previously inferred as 12, 14, and 9, respectively (Raven, 1975). Indeed, even in the presence of a strong phylogenetic signal (e.g., closely related species sharing similar chromosome numbers; Escudero *et al.*, 2012; Carta *et al.*, 2018), algebraic inferences of chromosome numbers become increasingly difficult with increasing phylogenetic depth, as identical chromosome numbers will occur in unrelated lineages (Weiss-Schneeweiss & Schneeweiss, 2013). The dataset analysed here is the most extensive ever used for inferring ancestral haploid number in angiosperms, but it still poses challenges concerning incomplete taxon sampling and phylogenetic resolution at the family level.

Our chromosome number evolution results support the conclusion that genome duplication and descending dysploidy were critical events in the evolution of angiosperms (Escudero *et al.*, 2014). Whilst polyploidization is the second most frequent transition type, ancient polyploidy events are underrepresented. The absence of polyploidization events at the base of the tree is in agreement with the maintenance of the ancestral haploid chromosome number *n* = 7 inferred in the deepest part of the phylogeny, with the inferred ancestral diploid status and with the very low ancestral genome size. Polyploidization events were instead inferred mainly toward the tips of the tree, partially supporting previous evidence revealing independent genome duplications near the base of several families (Escudero *et al.*, 2014; Leebens-Mack *et al.*, 2019; Cusimano *et al.,* 2012), leading to high haploid chromosome numbers. Indeed, this may be the case of many families in the Magnoliid clade. Our results do not support a link among some of the most extensive plant radiations and ancient polyploidization rounds (Jiao *et al.*, 2011). Nevertheless, the macroevolutionary dynamics of ploidy level changes here presented, emphasize the importance of diploidization events, suggesting that both WGD and PPD are widespread phenomena in angiosperms. Patterns of early WGD were suggested also in gymnosperms (Li *et al.*, 2015), where however polyploidy is rare (Rastogi & Ohri, 2020). On the contrary, repeated WGD events in the ancestors of monilophytes have contributed to their diversity and high chromosome numbers (Clark *et al.*, 2016). Inferring ancient polyploidy events from cytological data is a challenging task, because diploidization events (Schubert & Lysak, 2011; Mandakova & Lysak, 2018) following polyploidisation gradually can hide signals of genome duplication over time (Mayrose *et al.*, 2009). Yet, in our analyses, descending dysploidy was indeed the most common cytogenetic mechanism of chromosome number change, and this phenomenon is often regarded as a part of the diploidization process (Mandakova & Lysak, 2018). Our results on chromosome number evolution do not support WGD events at the origin of angiosperms or before it (Ruprecht *et al.*, 2017), but rather highlight the importance of WGD events later in the evolution of angiosperms. This is further corroborated by our results on genome size evolution, which clearly indicate that a) the ancestral genome size was small (1C = 1.73 pg, a value slightly higher than the inference by Leitch *et al.*, 2005) and mostly stable in the basal part of the tree; b) WGD likely did not occur in angiosperms for the first 30 million years of their evolutionary history (Fig. 3); and c) the majority of angiosperm nodes (57%) underwent a general genome size reduction.

Reconstructing the ancestral chromosome number is difficult, because there are no suitable outgroups for direct comparison (Doyle, 2012), and because extant early branching angiosperms (e.g., *Amborella* and Nymphaeales) are not necessarily holding plesiomorphic character-states. With these limitations in mind, we made inferences based on the distribution of *n* and genome size in extant angiosperms, and using probabilistic models accounting for various types of chromosome number transitions. Our study is not able to address the origin of either the chromosome number or genome size of the first angiosperms. Instead, it provides novel, detailed, and well-supported inference of ancestral haploid chromosome number and genome size of the common ancestor of all extant angiosperms. Interestingly, our inferred ancestral state for the haploid number *n* coincides with the ancestral basic chromosome number *p* previously proposed for angiosperms based on empirical counts (Ehrendorfer *et al.*, 1968; Stebbins, 1971; Walker 1972; Raven, 1975; Grant, 1981; Soltis *et al.*, 2005) or paleogenomic approaches (Salse *et al.*, 2012). In gymnosperms, the hypothesised ancestral number *n* = 12 (Nystedt *et al.*, 2013), based on the most frequent chromosome number in extant species, is partly supported by the reconstructed ancestral number in *Juniperus, n* = 11 (Farhat *et al.*, 2019). Not surprisingly, a higher ancestral chromosome number has been reconstructed for monilophytes, namely *n* = 22 (Clark *et al.,* 2016). According to Leitch *et al.* (2005) both gymnosperms and monilophytes showed an “intermediate” ancestral genome size (1C values ranging between 3.5 and 14.0 pg). Altogether, it appears that spermatophytes may have a lower ancestral chromosome number, compared to monilophytes, suggesting differences between the genomic evolutionary trajectories of these groups (Clark *et al.*, 2016).

Our study allowed to clarify a long-standing question (Stebbins, 1971), but such reconstruction necessarily comes with limitations. We do believe our results allowed a major step forward in understanding the ancestral chromosome number for angiosperms, and we believe that this issue should be added to the angiosperm macroevolutionary agenda (Sauquet & Magallón, 2018). Further progress in reconstructing the ancestral genome organisation in angiosperms may require the development of models that include heterogeneity in the patterns of chromosome evolution across a phylogenetic tree (Zenil-Ferguson *et al.*, 2018), along with a deeper insight into genome and karyotype evolution.

## Acknowledgements

The authors thank Marcial Escudero and Itay Mayrose for their help with ChromEvol analyses. We are also grateful to Rosana Zenil-Ferguson for her help with the chromploid package.

## Author contributions

A.C. planned and designed the research, analysed the data and wrote the manuscript. G.B. assisted in chromosome numbers acquisition. L.P. and G.B. contributed to successive versions of the manuscript and in solving theoretical and nomenclatural issues. All authors read and approved the final manuscript.

## Data Availability

All data analysed during this study are included in this published article and its supplementary information.

